# Towards personalized computer simulation of breast cancer treatment: a multi-scale pharmacokinetic and pharmacodynamic model informed by multi-type patient data

**DOI:** 10.1101/371369

**Authors:** Xiaoran Lai, Oliver M Geier, Thomas Fleischer, Øystein Garred, Elin Borgen, Simon Wolfgang Funke, Surendra Kumar, Marie Elisabeth Rognes, Therese Seierstad, Anne-Lise Børresen-Dale, Vessela N. Kristensen, Olav Engebraaten, Alvaro Köhn-Luque, Arnoldo Frigessi

**Author notes:** Joint last authors (Alvaro Köhn-Luque, Arnoldo Frigessi).

## Abstract

Mathematical modeling and simulation have emerged as a potentially powerful, time and cost effective approach to personalized cancer treatment. The usefulness of mechanistic models to disentangle complex multi-scale cancer processes such as treatment response has been widely acknowledged. However, a major barrier for multi-scale models to predict the outcomes of therapeutic regimens in a particular patient lies in their initialization and parameterization which need to reflect individual cancer characteristics accurately. In this study we use multi-type routinely acquired measurements on a single breast tumor, including histopathology, magnetic resonance imaging, and molecular profiling to personalize parts of a complex multi-scale model of breast cancer treated with chemotherapeutic and anti-angiogenic agents. We model the dynamics of drugs in tissue (pharmacokinetics) and the corresponding effects on their targets (pharmacodynamics). We developed a open-source computer program that simulates cross-sections of tumors under 12-week therapy regimes and use it to individually reproduce and elucidate treatment outcomes of four patients. For two of the tumors that did not respond to therapy, we used model simulations to suggest alternative regimes, depending on their individual characteristics, with improved outcomes. We found that more frequent doses of chemothereapy reduce tumor burden in a low proliferative tumor while lower doses of anti-angiogenic agents improve drug penetration in a poorly perfused tumor. In addition to bridge multi-type clinical data to shed light on individual treatment outcomes, our approach identified a few tumor-related aspects that need to be clinically portraited better to allow for future model-driven personalized cancer therapy.

## Introduction

Current personalized cancer treatment is based on a few biomarkers which allow assigning each patient to a subtype of the disease, for which treatment has been established [22]. In breast cancer, multigene tests like Mammaprint or PAM50 give prognostic information to guide clinical decisions [15, 27]. Such stratified patient treatments represent a first important step away from one-size-fits-all treatment. However, the accuracy of disease classification comes short in the granularity of the personalization: it assigns patients to one of a few classes, within which heterogeneity in response to therapy is still large [36]. In each of these classes, randomized clinical trials (RTC) can be run, in order to compare a few treatment regimens and to identify the best on-average one. As there is a combinatorial explosive quantity of combinations of cancer drugs, doses and regimens, only very few can be explored by RCTs. The concept of ‘one disease in each patient’ challenges diagnosis and therapy.

Mechanistic mathematical modeling and simulation have emerged as a powerful approach to investigate the influence of biological factors on tumor progression and therapy response [2– 4]. Current models are able to account for complex interactions at the cellular and molecular level, and are capable of bridging multiple spatial and temporal scales in ways that would be impossible using experimentation [9, 19]. Successful multi-scale models can describe with acceptable approximation the dynamics of tumors under the effect of a specific therapy. The present computing capacity allows exploration of a large number of treatment regimens by running multiple model simulations in parallel. Yet, considerable challenges hinder the use of computational modeling to guide patient’s treatment in clinical practice [4, 10]. The choice of the level of biological detail represented in the mathematical model can always be questioned and a judicious balance between biological and model complexity has to be found. An essential question is whether at all mathematical models can be personalized to predict the effect of therapies in each specific patient [4, 34]. This requires individualized initialization and parameterization of such a model which are typically difficult to perform.

For this paper, our team of oncologists, pathologists, molecular biologists, medical imaging physicists, statisticians and mathematicians examined whether a computational modeling approach can be used to effectively simulate individual outcomes of a specific group of patients with invasive breast cancer receiving different treatment regimes (described later). We asked the following questions: Is it possible to design a mathematical model that integrates individual patient’s data collected in routine clinical practice to reproduce patients’ outcome? Can such a model be flexible enough to simulate a wide range of possible responses by capturing fundamental biological processes at an acceptable level of approximation? Are the available data sufficient to personalize such a mathematical model? This paper answers positively the first two questions: we developed a model which allows *in-silico* simulation of the treatment outcome. Regarding the third question, we show the need to measure more precisely certain individual drug and tumor characteristics, in order to estimate certain parameters which modulate the dynamics.

Barbolosi et al. [4] underline the importance of capturing in a mathematical model pharmacokinetics (the movement of drugs in the biological tissues) and pharmacodynamics (their mechanisms of action and effect). Both are present in our model. Furthermore Barbolosi et al. [4] distinguishes between phenomenological (descriptive) models and mechanistic (explicative) models. Phenomenological models are parametric models of the time evolution of the tumor that do not rest on explicit biological mechanisms. Mechanistic models instead, account for specific biological details and dynamics. We present here a mechanistic model that captures non-linear, multi-scale dynamics in space and time. We consider discrete individual cells and blood vessels together with continuous quantities like oxygen pressure and drug concentrations; such models are termed ‘hybrid’. Gallo and Birtwistle [13] use the term ‘enhanced pharmacodynamics’ for models that merge multi-scale dynamics with pharmacokinetics, as is in our case. An important distinction is between models informed by individual patient data and those who are not, and therefore cannot be used for personalized treatment optimization. While there are many studies in the literature which propose models in some of these dimensions (for instance [1, 6, 8, 16, 21, 23, 26, 28–30, 33, 35]), our study is possibly the first one towards personalized computer simulation of breast cancer treatment incorporating relevant biologically-specific mechanisms and multi-type individual patient data in a mechanistic and multi-scale manner.

We inform our model with data from four breast tumors collected in a recent neoadjuvant clinical phase II trial [32]. Patients included in this study were randomized in two arms to receive chemotherapy with or without bevacizumab. Histological, magnetic resonance imaging (MRI) and molecular data were collected before, during and at the end of neoadjuvant treatment. We develop precise pipelines for clinical data preprocessing, model initialization and personalization. Besides individual histological and MRI data, our model also makes use of some genomic patient data. By means of extensive numerical simulations, we show, as a proof-of-concept, that patient-specific and multi-scale modelling allows us to reproduce treatment outcomes of the four patients. In addition we investigate if and how alternative treatment protocols would have produced different outcomes. This is a first step towards virtual treatment comparison. Finally, we suggest which additional data and experimentation with tumor material is needed in order to improve the accuracy of the simulation, towards precise outcome prediction. Our study shows that simulation-based personal treatment optimization is feasible and powerful, and should be developed further as a promising avenue of personalized treatment.

## Materials and methods

### Patients and treatment

We selected four patients with HER-2 negative mammary carcinomas from the NeoAva cohort [32], a randomized, phase II clinical trial that evaluated the effect of bevacizumab in combination with neoadjuvant treatment regimes during 24 weeks. See section S4.1 in Supplementary Material (SM) for more details. We analyse only the first 12 weeks, where patients were treated with chemotherapy (FEC100, 4 courses) and randomized 1:1 to receive bevacizumab (15 mg/kg every third week given concurrently with chemotherapy) or not. The four patients had either a complete or no response by MRI at 12 weeks of treatment. An overview of baseline characteristics, treatment and response for the four patients is shown in table 1.

**Table 1:** Patient overview. Baseline characteristics are shown together with assigned clinical trial arms and tumor volumes measured from DW-MRI segmentation at weeks 0, 1 and 12 after initiation of the treatment. Volumes marked with asterisks were unavailable due to problems with fat suppression and instead computed from Dynamic Contrast Enhanced (DCE) segmentation. As patient 1 was treated with bevacizumab, typically reducing DCE-MRI signal, maximum tumor diameters were also used (30mm at week 0 and 31mm at week 12). Putting this together, at week 12, patients 1 and 3 were classified as non-responders (NR) while patients 2 and 4 were classified as complete responders (CR). † Invasive Ductal Carcinoma ‡ A few possibly malignant cells not further classified were seen in lymph node aspirate before treatment start.

| Patient ID |  | 1 | 2 | 3 | 4 |
| --- | --- | --- | --- | --- | --- |
| Age |  | 48 | 31 | 40 | 41 |
| Histological tumor type |  | IDC<sup>†</sup> | IDC | IDC | IDC |
| Histological tumor grade |  | 2 | 3 | 3 | 2 - 3 |
| Hormone Receptor status | ER | positive (100%) | positive (1%) | negative | negative |
|  | PR | positive | negative | negative | positive |
| Lymph Node (LN) status | Week 0 | positive | negative | positive | negative<sup>‡</sup> |
| PAM50 subtype |  | luminal B | basal | HER2 | basal |
| Clinical Trial Arm |  | FEC100 + bevacizumab | FEC100 + bevacizumab | FEC100 only | FEC100 only |
| Tumor volume ($mm^3$) | Week 0 | 3987 | 14940 | 11449.5* | 1934 |
|  | Week 1 | 3392 | - | 11460 | 1279 |
|  | Week 12 | 1662* | 89 | 11780 | 0 |
| Response at week 12 |  | NR | CR | NR | CR |

### Histopathology

The histopathological analysis was performed on needle biopsies taken from the breast tumors at week 0. See details and used images in section S4.2 of SM. Fiji and its cell counter plugin [31] were used for manual identification of cancer and stroma cells in each image.

### Magnetic resonance imaging

Patients were examined on a 1.5 T MRI scanner (ESPREE, Siemens, Erlangen, Germany) at weeks 0, 1 and 12 of the neoadjuvant treatment. See section S4.3 in SM for a description of the imaging protocol and data, including dynamic contrast-enhanced (DCE-MRI) and diffusion weighted (DW-MRI) imaging. For DCE-MRI analysis, an extended Tofts model was used, which included the determination of the contrast-enhancement curve of the contrast agent in each individual voxel (volume 1 mm × 1 mm × 1.5 mm). DW-MRI data were analyzed using a simplified IVIM model-based analysis as described in [7].

### Molecular data

To estimate parameters representing subcellular processes, we assessed a number of molecular features for each tumor. mRNA levels of VEGFA and *TP53* in the tumor samples were determined using one color SurePrint G3 HumanGE 8 60 k Microarrays (Agilent Technologies) as described in [32], available in the ArrayExpress database, accession number E-MTAB-4439. The PAM50 subtyping [25] was used to assign a subtype to each sample in the NeoAva cohort and proliferation score was derived for each tumor by computing mean expression values of the 11 proliferation-related PAM50 genes [25, 32]. *TP53* mutation status was determined by sequencing the entire coding region (exons 2-11), including splice junctions as described in [32]. Furthermore, pathway deregulation score (PDS) [11] of the Hypoxia-inducible factor 1-alpha (HIF1A) pathway was calculated for all four patients at screening and at week 12 and normalized against 50 tumor-free samples.

### Mathematical model

To model the response of a cross section of tumor tissue to a combination of chemotherapeutic and antiangiogenic drugs, we use a hybrid cellular automaton model, describing multi-scale pharmacokinetic and pharmacodynamic processes relevant to the therapy response. We combine cellular, extracellular and intracellular dynamics and inform them by multi-type, individual patient data. The model is fully described in section S1 of SM. Briefly, we describe individual cells and cross sections of functional blood vessels in a 2D section of tumor tissue as discrete agents on a regular grid. Cell division and death as well as blood vessel formation and removal, are controlled in the cellular automata by intracellular and environmental factors, described by ordinary and partial differential equations. For instance, the concentration of each drug in the blood and in the tumor tissue is described by ordinary (pharmacokinetics) and partial (reaction-diffusion) differential equations respectively. As FEC chemotherapeutic agents can only kill cycling cells, we consider a simple cell cycle model for each cell and model the effect of low oxygen tension as delaying cell division. We account in each cell for the effect of hypoxia to *TP53* and VEGF expression using a system of ordinary differential equations. This allows us to model the amount of VEGF produced by each cell and the inhibition of VEGF molecules by the antiangiogenic drug. To model the effect of VEGF on the tumor vasculature, we use a stochastic model where the probability of formation or removal of vessels is influenced by the local VEGF concentration.

The workflow to run a personalized simulation of drug response of a breast tumor portion is outlined in fig. 1. Tumor screening data at week 0 are used, as described in section S2 of SM, to initialize and parameterize the model, while MRI examinations at weeks 1 and 12 are used to validate the simulated outcomes (box A in fig. 1). After initializing and parameterizing the mathematical model using patient data and public data in literature (boxes B and C), we simulate the drug schedule used in the clinical trial for each patient (box D). For that, we increase the amount of the corresponding drugs (FEC100 +/-bevacizumab) in the blood at the time points of administration according to the dosage. Each computer simulation then runs a complete cycle of 12 weeks of the spatio-temporal dynamics of the considered tumor portion under the effect of the applied drugs (box E). Computational details and a link to the open-source computer code can be found in section S3 of SM. Simulated outcomes include the spatial distribution of cells, blood vessels and all considered molecules at any time between the start of the therapy and week 12 and in particular at the end of the study period (box F). To validate a simulation, we compare the simulated outcomes with the actual ones (box H). For that we use clinical tumor volumes and apparent diffusion coefficients (ADC) [32] calculated from DW-MR imaging at week 1 and 12 (box G). As we are simulating only very small portions of the tumor bulk, we selected only patients that were complete responders (number of tumor cells in all simulated tumor portions at week 12 should be around zero) or complete non-responders at week 12 (approximately the same number of tumor cells in week 12 as in week 0). Moreover, changes in ADC histograms of the segmented tumor volume at weeks 0, 1 and 12, which relate to changes in the tumor cellularity [33], should qualitatively agree with the number of tumor cells in our simulations.

**Figure 1:**
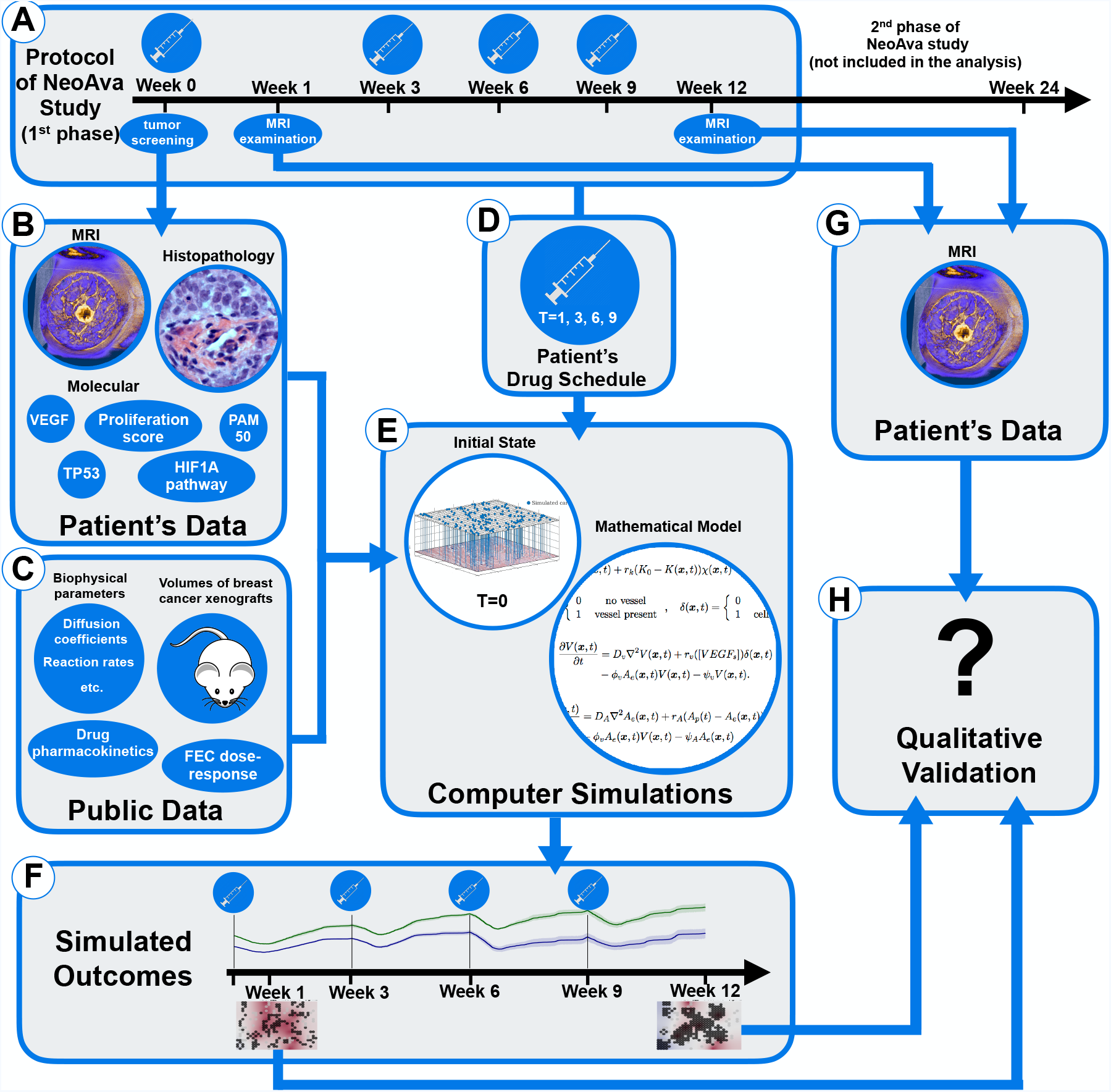
Simulation and validation workflow, outlined through panels A to H

## Results

### Comparison of patient characteristics and treatment responses

We consider four patients numbered from 1 to 4, see table 1 for an overview of all patients and section S4 of the SM for the complete clinical data set we used. Patients 1 and 2 received identical therapy (FEC100 plus bevacizumab) with the same dose and schedule. Patient 1 did not respond to the therapy after 12 weeks, while patient 2 responded well. At baseline, patient 2 had a higher tumor cell density and larger tumor size than patient 1, as shown by histological images and DW-MRI. This coincided with higher PDS of HIF1A pathway at the start of the treatment. Their tumors were also classified differently as Basal-like and Luminal B, respectively, and the estimated proliferative capacity of patient 2 was much higher than that of patient 1 (see parameter *T_min_* in section S2.5.1 of the SM). This is in agreement with the estimated PAM50 proliferation scores [24] of both patients. Other molecular and MRI-deduced parameters used in this study, did not differ significantly. Specifically, expression levels of VEGF and *TP53* were almost identical, and both tumors were *TP53* wild-type. The typical values of perfusion parameters *k*_trans_ and *υ_p_* estimated from DCE-MRI, were similar too.

Patients 3 and 4 received only FEC100 with the same dose and schedule. Patient 3 did not respond to the therapy after 12 weeks, while patient 4 responded completely. The tumors were classified as HER2 enriched and Basal-like respectively, but the patients had very similar proliferative capacity as shown by their estimated PAM50 proliferation scores. They also differed in tumor morphology, vessel perfusion and *TP53* status and expression. Comparing histological slices and DW-MRI, the tissue of patient 3 was densely packed with cells, while patient 4 showed more heterogeneity. DW-MRI of patient 4 also showed heterogeneity. At week 0, DCE-MRI showed that the tumor in patient 3 had a very poorly perfused core while it was highly perfused on the outer edge, resulting in cross-sections with ring-like patterns. Interestingly, DW-MRI analysis suggests that the tumor core is not necrotic. The tumor from patient 3 was *TP53* wild-type while the tumor from patient 4 was *TP53* -mutated. Expression levels of *TP53* were higher in patient 4 than in patient 3 while VEGF expression was comparable and very high in both patients. Patient 3 also had much higher HIF1A PDS compared to Patient 4 at week 0, exhibiting signs of a denser and more hypoxic tumor.

### Model simulations reproduce treatment outcomes for all patients

For each of the four patients we run personalized simulations of tumor portions under the treatment received in the clinical trial. We then use the response data obtained from MRI at weeks 1 and 12 to compare the simulated and actual outcomes. The simulation and validation workflot is outlined in fig. 1 and described in the SM.

Model simulations of the outcomes of patient 1 and patient 2 are shown in fig. 2. We used two initial biopsy portions and corresponding configurations of cancer and stroma cells for each patient (Biopsy A and B). For each such configuration, we ran ten independent stochastic simulations of the 12-week treatment period using the same initialization and parameterization, but different random events such as births and deaths of vessels. We plot the time evolution of the cancer cell density, defined as the proportion of grid points occupied by cancer cells at any given time, for the ten simulations. Simulated cell densities for patient 1, shown in fig. 2a, decrease moderately after drug administration and grow in the period between consecutive administrations. The four grid snapshots show, at different time points, the spatial distribution of cancer and stroma cells together with the oxygen level in one representative simulation. All of the twenty simulated experiments presented moderate degrees of hypoxia, as seen by the white and light blue background displayed in the snapshots of fig. 2a. This is in agreement with the observed hypoxic pathway activity and VEGF expression, shown in fig. S18b and fig. S18d of the SM. In fact, despite of production of VEGF by hypoxic cells, the applied dose of bevacizumab reduces VEGF concentration in the tumor tissue to very low levels, as recorded above each snapshot in fig. 2a. In our model blood vessels disappear with a certain probability under low VEGF concentration. For patient 1, if this probability is high enough, so that vessels tend to be absent, the tumor can persist, because bevacizumab reduces tumor perfusion and the chemotherapeutic agents cannot reach the tumor tissue. We can reproduce the outcome at week 12 using any value for the probability for vessels to become dysfunctional under low VEGF and an intermediate value of the chemosentisivity parameter, as shown in fig. S3 and fig. S6 of SM. In fig. 2a, even when using a very low probability of vessel disappearing (*p*_death_ = 0.0001), the killing of cancer cells after each dose administration does not prevent cancer cell to proliferate again by the next drug administration, resulting in a net balance between killing and proliferation in 20 out of 20 simulations. This is in agreement with the available ADC data of patient 1 at week 0 and 1, showing little difference in tumor density (fig. S17a of SM). By the end of the simulations, the cancer cell density is approximately the same as it was at the beginning.

**Figure 2:**
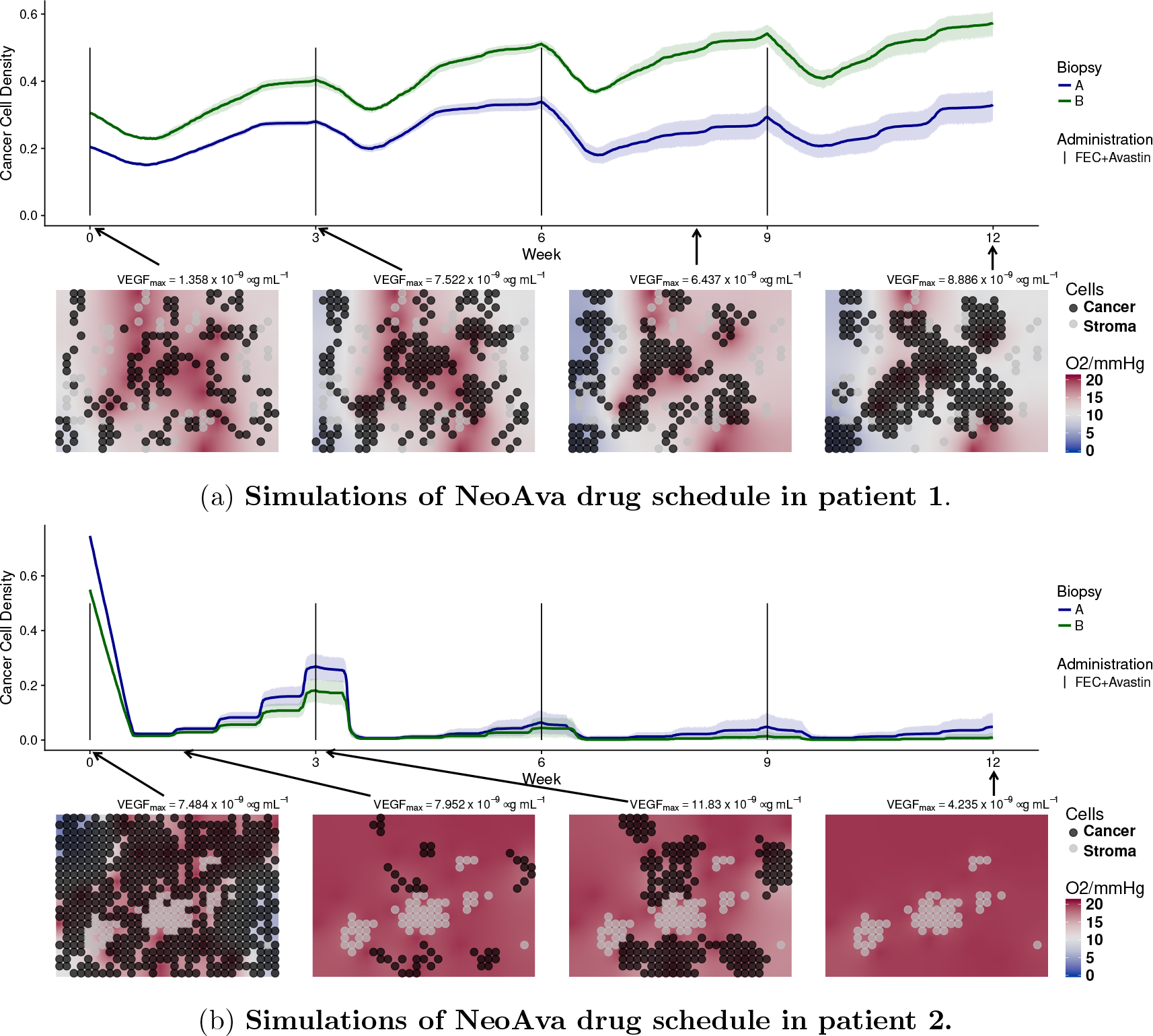
Simulated time evolution of the cancer cell density starting from two different cell configurations at week 0 corresponding to biopsy portions A and B. Each line represents the average cancer cell density of 10 independent stochastic simulations. The corresponding color band indicates the 95% bootstrap confidence interval. Lower panel of each figure shows the spatial distribution of cancer and stroma cells for a representative simulation of the biopsy portion B for each patient. Background color represents oxygen pressure in mmHg.

On the other hand, patient 2 is a responder. A significant number of blood vessels should remain in the tissue to allow enough chemotherapeutic drugs to be distributed and thus kill most of the tumor cells. To reproduce the available ADC data, where we see that virtually no tumor remained at week 12 (fig. S17b), the following is required. First, a low probability of vessel disappearing (*p*_death_ < 0.01), so that part of the tumor vasculature is resistant to the VEGF inhibitor) and, second, a chemosensitivity higher enough (*β* > 6000), so that not many cells escape from the therapy after each administration, see fig. S3 and fig. S6 of SM. In the simulations of patient 2 shown in (fig. 2b), all tumor cells were killed after 12 weeks in 19 out of 20 experiments. Due to higher cell densities at week 0, hypoxia is more severe than for patient 1, all in agreement with the available HIF pathway data (fig. S18d of SM) and ADC data (fig. S17b of SM).

Patient 3 is a non-responder. As revealed by MRI data (fig. S15 in the SM) the core of the tumor is poorly perfused, contrary to the border. We show in fig. 3 simulations for both perfusion profiles. We used the estimated values *k*_trans_ = 0.0067 min^−1^ and *υ_p_* = 1.73% that reflect poorly-perfused condition in the core, see fig. 3a. Since the number of vessels is small and their permeability is low, the simulated drug concentration arriving in the tissue is very low. As a result of this, free spaces in the simulation grid were taken over by the cancer cells, see fig. 3a. This is in agreement with DW-MRI data (fig. S17c) and HIF1A PDS which explain patient 3 being a non responder. For the tumor edge, we used the estimated values *k*_trans_ = 0.2145 min^*−*1^ and *υ_p_* = 15.35% that lead to a highly-perfused condition. In this case, drugs are arriving in the tissue more efficiently. Provided cell chemosensitivity is high enough, simulated cancer cell densities can be reduced, as shown in fig. 3b. Although cells in the edge can be killed, this area could have been repopulated by the highly-proliferative cells from the core. As we only simulate small portions of the core and of the edge and we do not considered cell migration in our model, we have not tested this possibility.

**Figure 3:**
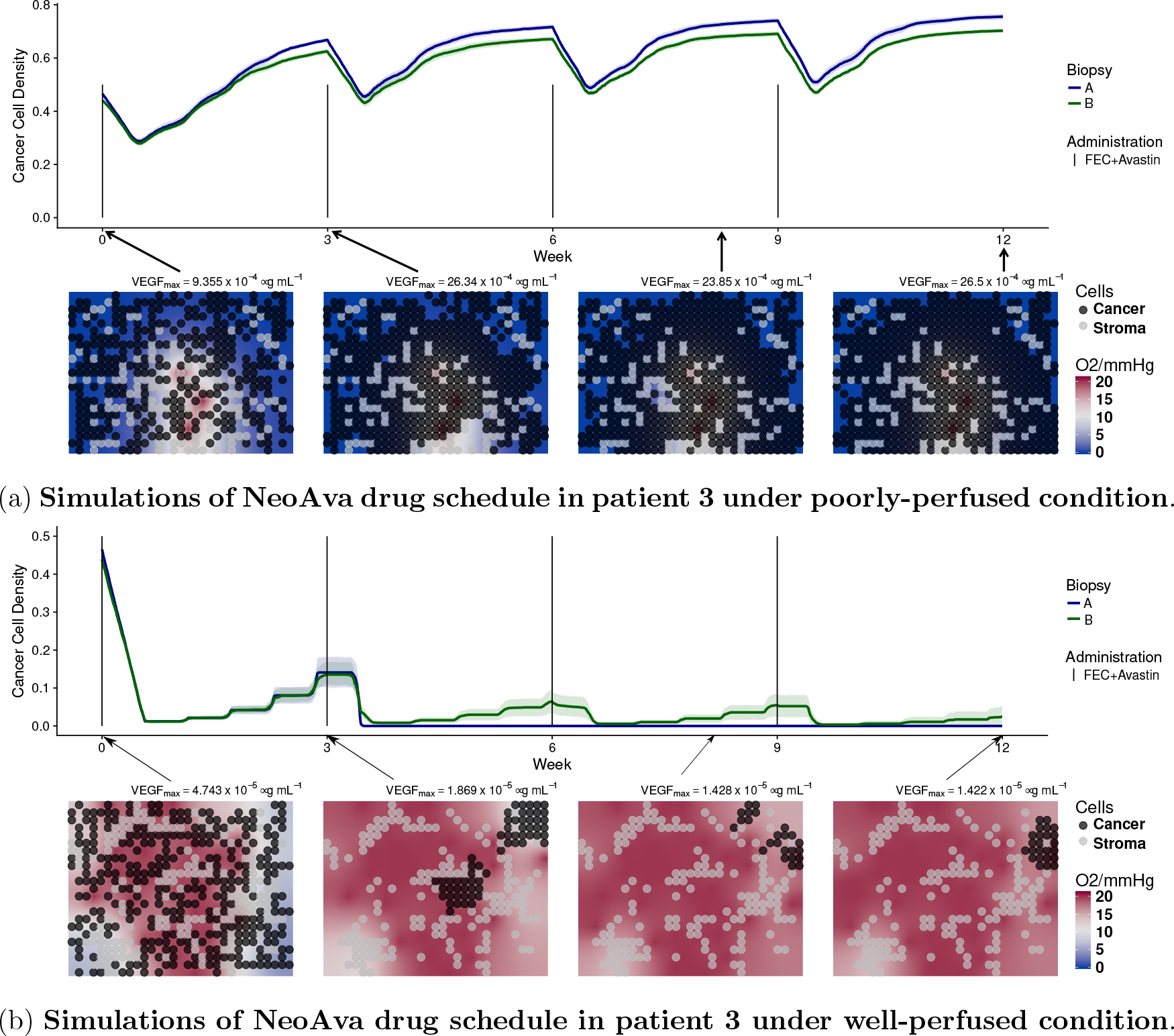
Time evolution of cancer cell density for two different cell configurations, biopsy portions A and B, and drug schedules under two different perfusion profiles. Each line represents the average cancer cell density of 10 independent stochastic simulations. The corresponding color band indicates the 95% bootstrap confidence interval. Figure 3a represents the core of the tumor of patient 3, while lower panel of fig. 3b shows the peripheral spatial distribution of cancer and stroma cells for a representative simulation of the biopsy portion B for each patient. Background color represents oxygen pressure in mmHg.

Patient 4 is a responder with also a heterogeneous perfusion profile as seen by DCE-MRI. But contrary to patient 3, a scattered pattern with two representative perfusion profiles is observed, one highly perfused with low permeability, the second with lower perfusion but high permeability. We show simulations of the first perfusion profile in fig. 4a, where we used the estimated parameters *υ_p_* = 6.0% and *k*_trans_ = 0.0067 min^*−*1^. Simulations of the second profile are shown in fig. 4b, where we used the estimated parameters *υ_p_* = 3.99% and *k*_trans_ = 0.1107 min^*−*1^. To reproduce the DW-MRI data in both cases, where we see that the tumor was drastically reduced at week 1 and almost disappeared by week 12 (fig. S17d), a higher rate of vessel creation compared to the other three patients is required (fig. S5b and fig. S4 in SM). This was achieved by increasing the model parameters *p*_birth_ and High_*V*_, as shown in table S3. This allowed to transport the drug efficiently to the tissue, even in the case where the permeability is very small. The simulated reduction of severe hypoxia to normoxia shown in the snapshots of fig. 4 is also in agreement with the observed significant decrease in HIF1A PDS comparing week 0 to week 12 (fig. S18d in SM).

**Figure 4:**
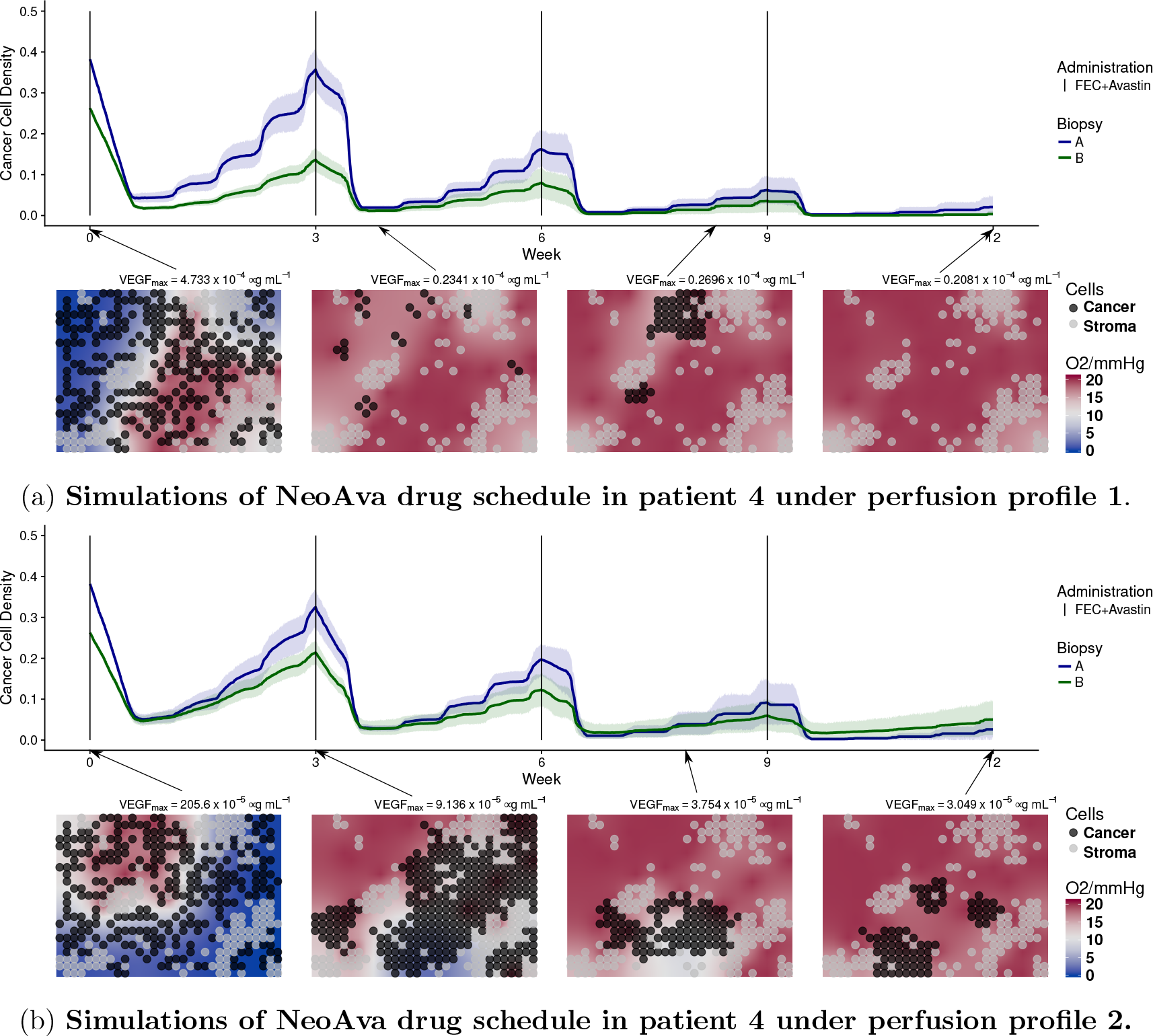
Time evolution of cancer cell density for two different cell configurations, biopsy portions A and B, and drug schedules under two different perfusion profiles. Each line represents the average cancer cell density of 10 independent stochastic simulations. The corresponding color band indicates the 95% bootstrap confidence interval. Lower panel of figs. 4a and 4b show the spatial distribution of cancer and stroma cells for a representative simulation of the biopsy portion B for the patient. Background color represents oxygen pressure in mmHg.

### Simulating alternative drug schedules

In addition to the regimes used in the clinical study, we also simulated alternative drug regimes (schedules and doses) for the two non-responders. The simulations of patient 1 in fig. 2a suggest that a main reason behind the survival of cancer cells after each drug administration was their low proliferation rate. As the interplay of treatment frequency and cell proliferation rate can contribute to the outcome, we hypothesized that administrating a smaller dose more frequently, while keeping the overall amount would be beneficial. We therefore simulated the following four chemotherapy regimes: every week with one third of the original dose, every one and a half week with half of the original dose, every two weeks with two thirds of the original dose and the original three week schedule used in the clinical trial. In fig. 5a we compare the outcomes after 12 weeks of treatment in terms of change in the initial cell density. We used the same initialisation and parameters of the personalised simulations of patient 1. Moreover, to see the effect of cell proliferation rate on the outcome, we repeated the same simulations but changed the cell cycle length parameter over a wide range. Simulation results show that the interplay between cell cycle length and schedule is complex and can exhibit non-monotonic behaviour. Interestingly, we see that drug administrations every week or every week and a half can improve the treatment outcome of patient 1 (marked with a star in fig. 5a). In fig. 5b, we show simulations of the most successful schedule obtained for patient 1, where we reduced FEC100 and bevacizumab to a third of its original dose, but administrated every week instead of every 3 weeks. By means of the higher drug administration frequency, the tissue was free of cancer cells after approximately 6 weeks. We also tested several regimens removing fluorouracil from the treatment. We found no difference in patient’s outcome comparing to the FEC100 regime administered every 3 week and every 2 week, agreeing with findings in clinical study [12], see fig. S19a in the SM.

**Figure 5:**
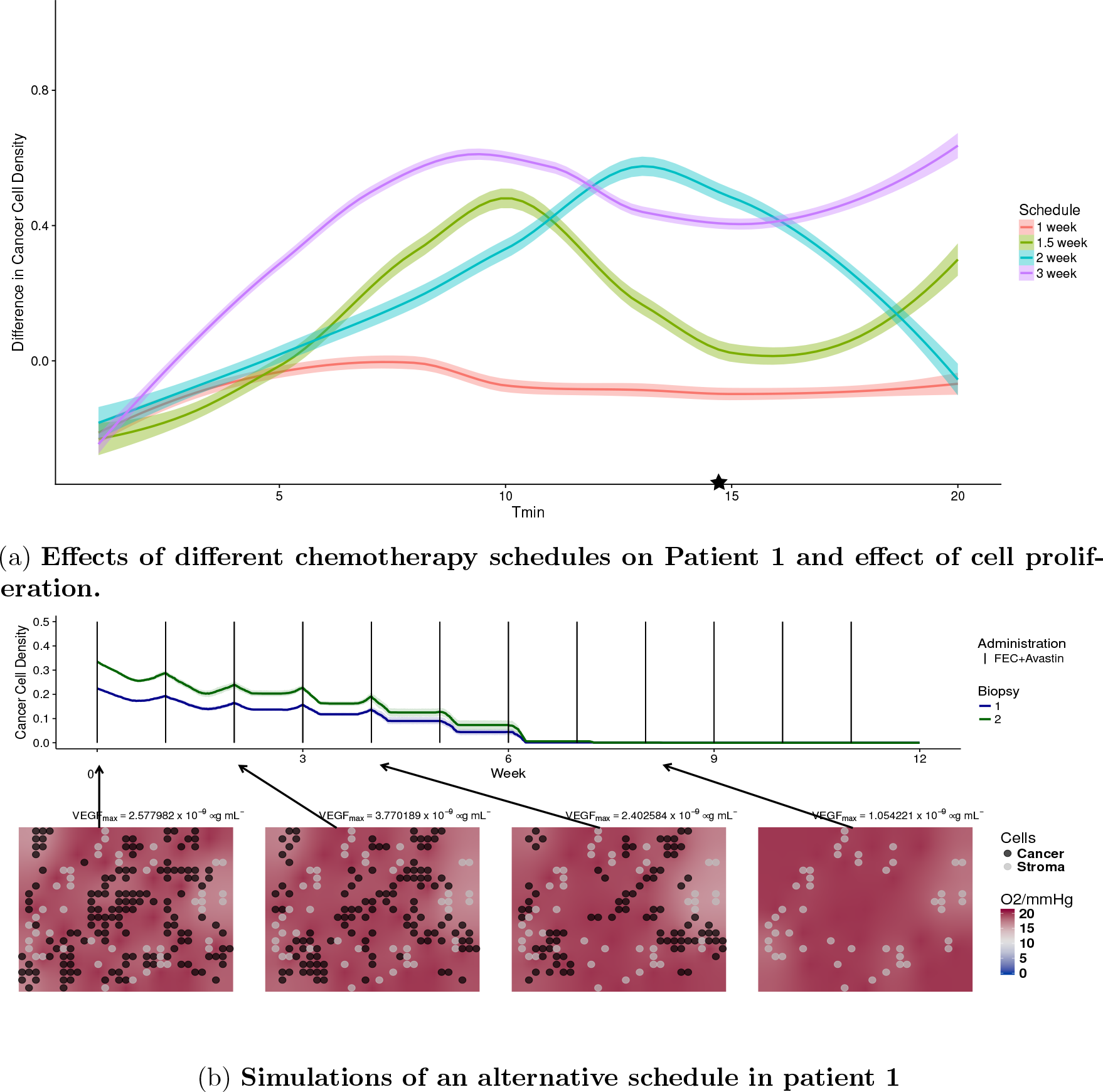
In fig. 5a, each colored dot represents the difference in cancer cell density between start and end of therapy under a specific drug schedule and minimal cell cycle length. *T*_min_ of daughter cells is randomised to introduce variability in cell cycle duration. Colored lines represent the locally weighted smoothed curve smoothing fitting of the simulations of different cell cycle length under the same drug schedule. The simulations of patient 1 corresponding to the value *T_min_* = 14.69 are labelled with a star. fig. 5b illustrates the temporal dynamics of the alternative drug schedule providing better outcome. Solid lines represent the average cell density (n=10) of the alternative therapy obtained by reducing FEC100 and bevacizumab to a third of its original dose, and administrating them every week instead of every third week.

For patient 3, who is a non-responder in the trial arm with chemotherapy only, our analysis suggests that the main reason for the negative outcome was that drugs do not penetrated enough in the tumor core. We hypothesized that by administrating an appropriate amount of bevacizumab, VEGF levels could be reduced appropriately, and tumor perfusion could be improved with the creation of new functional vessels in the core [17, 18]. Thereafter chemotherapy could be delivered efficiently and improve the outcome of the patient. We tested if tailoring a specific bevacizumab regimen to patient 3 would lead to better outcome. We added different bevacizumab regimens to the personalised simulations of patient 3, using the same initialisation and parameters. Moreover, to see the effect of VEGF expression on the outcome, we repeated the same simulations but changing the parameters modulating VEGF expression levels from low to high (fig. 6a). Since bevacizumab has a long half-life, the same schedule was used as for the chemotherapy (every three weeks) while scaling the amount with respect to what originally administered in the other arm of the trial. We see in fig. 6a that applying full amount of bevacizumab led to no improvement comparing to chemotherapy only, and that the optimal bevacizumab regime is different depending on the VEGF expression. In fig. 6b, we show simulations using the optimal bevacizumab regime for patient 3 (VEGF Level = 2.32). Comparing this simulation to its actual clinical outcome, the new therapy appears to improve the outcome by inhibiting the growth of the tumor, reducing the density of the tumor by 50% on average.

**Figure 6:**
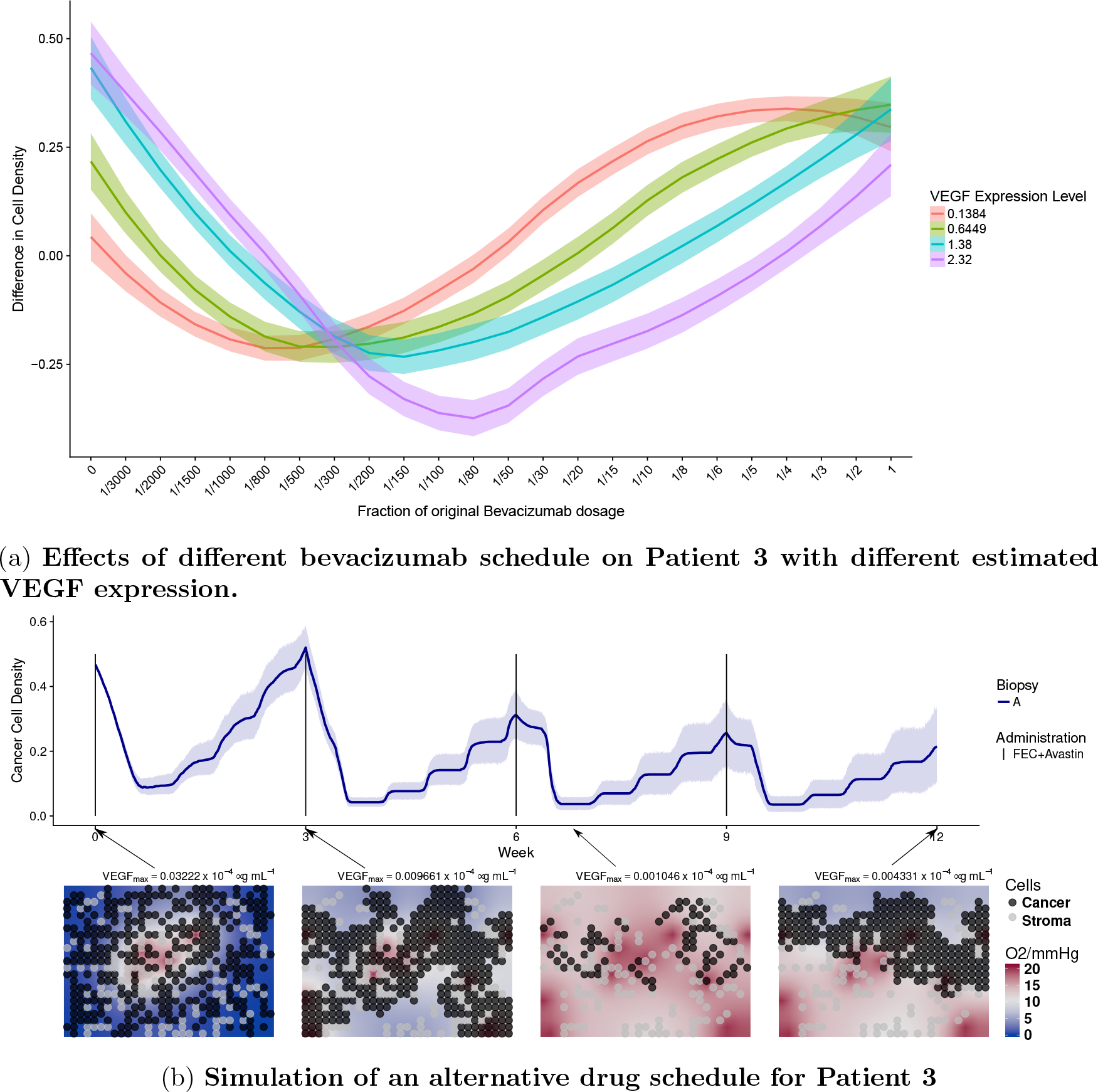
In fig. 6a, the y-axis represents the difference in cancer cell density between week 0 and week 12. Patients are treated under a 3-week administration interval. The x-axis indicates the quantity of bevacizumab administered to the patient as fractions with respect to the full dose, corresponding to 15 mg/kg. Each colored band summarizes 10 independent simulations for one patient with VEGF expression given in the legend. These four expression levels correspond to those observed in our four patients. However all other parameters are fixed as for Patient 4. In fig. 6b, we showed temporal dynamics of an alternative drug schedule providing better outcome. Solid lines represent the average cell density (n=10) of the alternative therapy obtained by administrating a reduced bevacizumab dosage of 0.5mg per kg body weight, equivalent of 3.33% of the dosage in the experimental arm along side of chemotherapy following a 3-week interval.

## Discussion

This paper is a new step towards the simulation of individualized tumor response to therapy. We have shown that it is possible to design a multi-scale mathematical model, which integrates different types of individual patients data collected in routine clinical practice and simulates the observed patient outcome. Our model is flexible enough to simulate very different outcomes based on individual data, by capturing fundamental biological processes at an acceptable level of approximation. Importantly, our model suggests possible mechanistic explanations of individual treatment outcomes and allows virtual testing of alternative treatment plans. The results are promising and potentially applicable to other solid cancers. We are now better equipped to recognise important limitations of this approach that need to be addressed in the future.

First, for each tumor under consideration we have only simulated a few sections of 200×300 microns, which might not be representative of the whole tumor bulk. Although the current computational power allows multi-scale model simulations of larger tumor portions, the approach is still limited by the availability of clinical data to inform them. For instance, clinical information at the cellular level, such as the number, position and type of cells, is nowadays possible only for biopsies. We are currently extending our algorithms to run cross-sections of full biopsies. A second limitation of our current model relates to tumor heterogeneity. Some cell clones can be more proliferative than others, produce more VEGF or be more resistant to therapy, for instance. Vessels can be different in size, have unequal functionality and permeability, contain different oxygen and drug concentrations, etc. Due to the current impossibility of characterizing to that level, cells and vessels heterogeneity from the available clinical data, our simulations assume all cells and vessels in a simulated tumor section to be of the same type. Extending our model to multiple clones, competing for resources would rely on a deeper picture of the patients’ tumor than routinely available today. Moreover, we have assumed that model parameters do not vary in time during the simulated treatment term. For instance we use constant drug chemosensitivity, which is a simplification and neglects the possible evolution of resistant phenotypes. In order to incorporate such details in the present approach, monitoring of tumor evolution is needed. We incorporated some degree of heterogeneity in our studies by running simulations with different cell configurations (observed in the biopsies), and different perfusion and permeabilities (observed in MR images of the whole tumor). We account for spatio-temporal heterogeneity in the simulated tissue section where cell and vessel numbers, as well as oxygen, VEGF and drug concentrations change in space and time. The mode of action of the VEGF depletion treatment was assumed to be a reduction in the number of functional capillaries, resulting in local reduced supply of oxygen and treatment to tissue. However, the effect may be more complex, related to the perfusion and leakage of the tumor influenced by the VEGF, as in [20, 32]. See also [16] for a different susceptibility of VEGF withdrawal dependent on the immune activation in the tumor. Future version of the model should incorporate at least first principles of such mechanisms.

An important contribution of our paper is the identification of the patient’s individual parameters which currently cannot be estimated precisely enough from the present clinical data: the cell-cycle length, the chemosensitivity of tumor cells and the sensitivity of vessels to the local VEGF concentration. In our feasibility study, we calibrated these parameters, within a range of realistic values, by choosing the ones which allows simulating a trajectory that leads to the true endpoint observed for the patient in the clinical trial. In this way we argue that if we could estimate these parameters, based on patient data, then the model could predict the outcome. Because we are using the outcome to determine the value of these parameters, our paper does not perform prediction of the outcome, but shows mere feasibility of such a prediction, assuming these critical parameters being correctly estimated. There are several ideas on how these parameters could be estimated. Cell-cycle length and drug chemosensitivities can be measured using ex-vivo patient material, while sensitivity of vessels could be perhaps estimated using in-vivo experiments on mice xenografts [5, 20]. Another possibility is to use Approximate Bayesian Computation, suitable for complex models like the present one, to estimate individual parameters from observing the first cycle of treatment. Preliminary results in this direction are promising [14]. Additional actions would be potentially useful in order to improve the accuracy of the simulated therapy: biopsies and MRI data should be matched in order to know exactly from where in the MR image the biopsy was taken. This would allow more accurate estimation of parameters obtained from MRIs, for instance the vessel permeability. Repeated MR images at different time points matched to each other [34] would allow us to relate tumor features such as volume, perfusion and cell density along therapy. Furthermore, instead of using average drug pharmacokinetic models, longitudinal measurements of drug concentrations in the blood could be easily performed to personalized them [4].

In this paper we use some genomic data made from the biopsy material, but much more can be done. It is known that mutations, gene expression and copy number variations portrait properties of tumor cells some of which may have implication on treatment success. For example, the PAM50 gene expression signature [25] allows classification of breast cancers tumors into five distinct classes in clinical use. As these genomic features modulate the efficacy of therapy, our study suggests that the key parameters in our model, in particular proliferative capacity and chemosensitivity of tumor cells, will depend on such genomic features of the patient.

In conclusion, this paper shows realistic possibilities for simulation guided personalized therapy and indicates where further research should focus to make this possible.

## Acknowledgements

The research leading to these results has received funding from the European Union Seventh Framework Programme (FP7-PEOPLE-2013-COFUND) under grant agreement number 609020 - Scientia Fellows. We also would like to acknowledge the help received from the Department for Research Computing at USIT, the University of Oslo IT-department.

